# Making sense of disorder: Investigating intrinsically disordered proteins in the tardigrade proteome via a computational approach

**DOI:** 10.1101/2022.01.29.478329

**Authors:** Nora E. Lowe, Roger L. Chang

## Abstract

Tardigrades, also known as water bears, are a phylum of microscopic metazoans with the extraordinary ability to endure environmental extremes. When threatened by suboptimal habitat conditions, these creatures enter a suspended animation-like state called cryptobiosis, in which metabolism is diminished, similar to hibernation. In this state, tardigrades benefit from enhanced extremotolerance, withstanding dehydration efficiently for years at a time in a type of cryptobiosis called anhydrobiosis. Recent studies have demonstrated that the tardigrade proteome is at the heart of cryptobiosis. Principally, intrinsically disordered proteins (IDPs) and tardigrade-specific intrinsically disordered proteins (TDPs) are known to help protect cell function in the absence of water. Importantly, TDPs have been successfully expressed in cells of other species experimentally, even protecting human tissue against stress *in vitro*. However, previous work has failed to address how to strategically identify TDPs in the tardigrade proteome holistically. The overarching purpose of this current study, consequently, was to generate a list of IDPs/TDPs associated with tardigrade cryptobiosis that are high-priority for further investigation. Firstly, a novel database containing 44,836 tardigrade proteins from 338 different species was constructed to consolidate and standardize publicly available data. Secondly, a support vector machine (SVM) was created to sort the newly constructed database entries on the binary basis of disorder (i.e., IDP versus non-IDP). Features of this model draw from disorder metrics and literature curation, correctly classifying 160 of the 171 training set proteins (~93.6%). Of the 5,415 putative IDPs/TDPs our SVM identified, we present 82 (30 having confident subclass prediction and 52 having experimental detection in previous studies). Subsequently, the role each protein might play in tardigrade resilience is discussed. By and large, this supervised machine learning classifier represents a promising new approach for identifying IDPs/TDPs, opening doors to harness the tardigrade’s remarkable faculties for biomaterial preservation, genetic engineering, astrobiological research, and ultimately, the benefit of humankind.

## Introduction

Subsects of life that can withstand environmental extremes have been the subject of longstanding scientific fascination. Understanding the proteomics of these organisms may aid us in harnessing their unique extremotolerance to develop biotechnologies conferring this resilience to other creatures. One key extremotolerant organism is the tardigrade, commonly known as the water bear (Figure 1).

**Figure 1.**
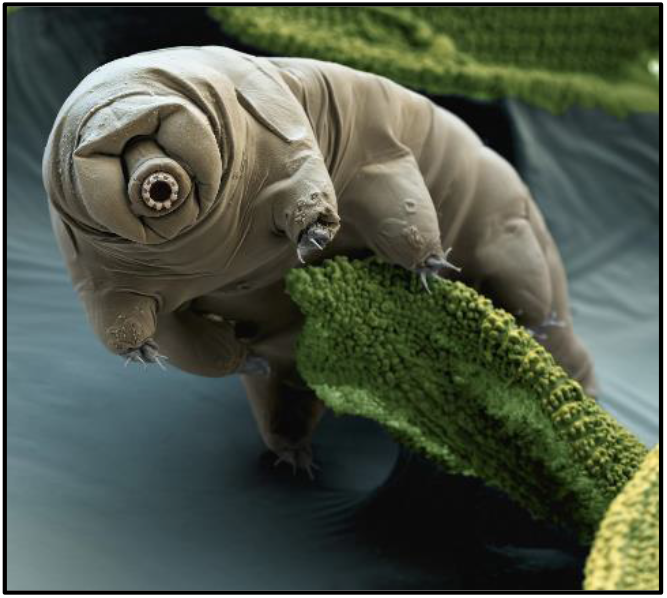
Scanning electron microscope image, with color enhancement, of a tardigrade specimen (National Geographic).

With their ability to survive environmental stresses sufficient to kill many other animals, the tardigrade has piqued the interest of researchers for 248 years (Bonnet & Goeze, 1773). These creatures are ubiquitous. There are over 1,300 species, all of which are meiofauna, minute animals inhabiting watery films and gaps between grains of sediment (Degma, Bertolani, & Guidetti, 2021). Tellingly, tardigrades have survived the five major mass extinctions; and Sloan, Alves Batista and Loeb (2017) even suggest it would require boiling Earth’s oceans to eliminate tardigrades definitively.

Understanding tardigrade extremotolerance is the central focus of this research. Currently, the literature includes robust documentation of the tardigrade’s capabilities in surviving environmental stresses, such as extreme temperatures (Doyère, 1842; Rahm, 1921), radiation (Beltrán-Pardo, Jönsson, Harms-Ringdahl, Haghdoost, & Wojcik, 2015; Hashimoto & Kunieda, 2017; Horikawa et al., 2006; Horikawa et al., 2013; Jönsson, Harms-Ringdahl, & Torudd, 2005), and vacuums/intense pressures (Jönsson, Rabbow, Schill, Harms-Ringdahl, & Rettberg, 2008; Seki & Toyoshima, 1998). In particular, tardigrades can withstand desiccation/dehydration adeptly; In 1948, zoologist Tina Franceschi described witnessing tardigrades from a 120-year-old dried moss sample being revived (Franceschi, 1948). Though this claim is heavily disputed (Jönsson & Bertolani, 2001), there is considerable contemporary evidence that tardigrades can indeed survive in a dehydrated state, also called *anhydrobiosis*, for nine to thirty years (Tsujimoto, Imura, & Kanda, 2016). This rare ability to cope with desiccation drives this computational investigation of the tardigrade proteome. Here, by pinpointing proteins involved in tardigrade desiccation tolerance, we provide insight for developing tardigrade protein-based technologies that could abate deleterious cell processes. This work could open doors for a range of applications, such as engineering drought and radiation-resistant plants, extending viability time for transfusion of blood products or transplant of organs, and establishing stable vaccine stockpiles.

Hence, in this study, we created and deployed a support vector machine (SVM) to generate a concise list of proteins of interest in tardigrade extremotolerance. Annotations of tardigrade-specific intrinsically disordered proteins (TDPs), a protein family recently implicated in tardigrade resilience (Hesgrove & Boothby, 2020; Yamaguchi et al., 2012), served as positive training data. We created and report here a novel, comprehensive, nonredundant pan-proteome database (PPD), from which we derived our training and testing sets. This database is composed of the phylum’s publicly available protein sequences, with flagged proteins involved in anhydrobiosis and other forms of extremotolerance. The feature set selected includes DISOPRED3 disorder metrics (Jones & Cozzetto, 2015), such as fractions of disordered residues and concentrations of certain amino acids, as well as phenotypic properties derived from literature curation. In short, this study consisted of creating an all-inclusive database and narrowing it down to locate proteins of interest. Altogether, understanding TDPs could help elucidate how to repurpose the tardigrade’s aptitude for survival for human benefit. This dataset, to the best of our knowledge, is the most comprehensive Tardigrada PPD that has been deduplicated to eliminate redundancy. An efficient regular expression-based algorithm permitted the assignment of a unifying identifier to most PPD proteins. Whereas previous proteomic studies have struggled with inconsistent nomenclature obfuscating protein identity, our database overcomes this obstacle. Overall, this study represents the first attempt of its kind to strategically mine proteomes for disordered proteins involved specifically in cryptobiosis. Ultimately, an enhanced understanding of such proteins could enable humans to imitate tardigrade resilience through development of novel TDP technology with wide-ranging applications in translational medicine and genetic engineering.

## Review of Literature

### Mechanisms of Anhydrobiosis

The tardigrade’s unique ability to undergo anhydrobiosis has been long documented but only recently understood. Anhydrobiosis is a type of cryptobiosis that can occur in any developmental stage (Schill & Fritz, 2008) whereby the tardigrade retracts its limbs and curls into a spherical formation called a tun (Figure 2, overleaf; Baumann, 1922). In a 1997 landmark study, Ricci and Pagani proposed a “Sleeping Beauty” hypothesis of aging, postulating that the tardigrade’s biological clock pauses during the tun state. Hengherr, Brümmer, and Schill (2008a) corroborated this idea by showing how lifespans of periodically dried *Milnesium tardigradum* were similar to those of their control counterparts, excluding time spent in the tun state. In this state, a type of *biostasis*, tardigrades rely on a host of molecular mechanisms to survive prolonged periods of desiccation, such as the disaccharide trehalose.

**Figure 2.**
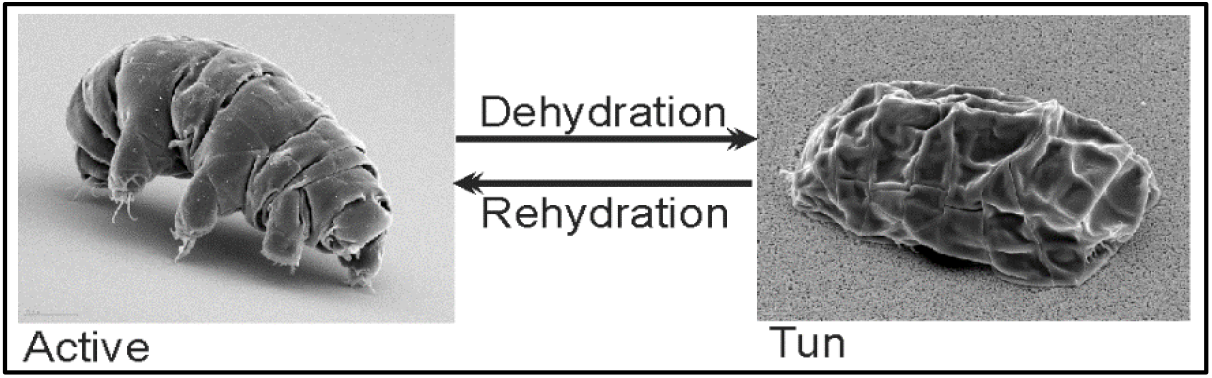
A comparison between active (left panel) and desiccated (right panel) states in tardigrades (Schokraie et al., 2010).

#### Implication of Trehalose

Trehalose is a sugar reported to be involved in anhydrobiosis in some tardigrade species (Crowe, 2002; Kinchin, 2008; Webb, 1964). However, a preponderance of inconsistencies, both between and within individual studies, reveal there is considerable inter-/intraspecies variation in concentrations of this substance. To illustrate, Jönsson and Persson (2010) observed increased levels of trehalose in *Macrobiotus islandicus* (accounting for 2.9% of anhydrobiote dry weight) compared to lower amounts in other species such as *M. tardigradum* (0.077% of dry weight). By contrast, in a different study, the latter species was previously shown to lack trehalose altogether (Hengherr, Heyer, Köhler, & Schill, 2008b). In tandem, these conflicting findings indicate trehalose many not be solely responsible for the phenomenon of anhydrobiosis in tardigrades.

While historically the field has spent a great deal of time focused on trehalose, scientists have recently redirected their attention to the tardigrade proteome. Promising findings have implicated proteins such as late embryogenesis abundant proteins (LEAs; Förster et al., 2009; Förster et al., 2012; Schokraie et al., 2010; Tanaka et al., 2015), heat shock proteins (Hsps; Alterio, Guidetti, Boschini, & Rebecchi, 2012; Förster et al., 2009; Förster et al., 2012; Jönsson & Schill, 2007; Reuner et al., 2010; Rizzo et al., 2010; Schill, Steinbrück, & Köhler, 2004; Schokraie et al., 2010; Schokraie et al., 2011; Wang, Grohme, Mali, Schill, & Frohme, 2014; Yoshida et al., 2017), and DNA damage suppressor proteins (Dsup; Hashimoto et al., 2016; Hashimoto & Kunieda, 2017; Yoshida et al., 2017) in augmenting tardigrade extremotolerance. Consequently, the tardigrade proteome has taken center stage as a cache for previously unexplored factors with ties to desiccation tolerance. In particular, proteins lacking order, meaning they do not have fixed tertiary structures, are of heightened interest.

### Tardigrade-Specific Intrinsically Disordered Proteins

A family of proteins called *tardigrade-specific intrinsically disordered proteins* (TDPs) is heavily involved in tardigrade cryptobiosis. Some TDPs are heat soluble, including cytosolic abundant heat soluble (CAHS), secretory abundant heat soluble (SAHS), and mitochondrial abundant heat soluble (MAHS; Yamaguchi et al., 2012). In addition, Dsup is type of nucleosome-binding and DNA-protecting protein (Chavez, Cruz-Becerra, Fei, Kassavetis, & Kadonaga, 2019) that is also considered a TDP. The term “tardigrade-specific” indicates that these proteins, to date, have not been identified outside of the phylum, and “intrinsically disordered” signifies proteins (IDPs) or protein regions (IDRs) lacking a consistent tertiary structure under certain cellular conditions (Jirgensons, 1996). In essence, unique properties of the TDP family can be attributed to this disorder, in that proteins within this classification adopt a material state upon desiccation, forming non-crystalline, amorphous solids (Boothby et al., 2017). Considered a type of biological glass, these solids have been shown to protect cells during dehydration in a multitude of ways (Crowe, Carpenter, & Crowe, 1998; Sun & Leopold, 1997) by coming together to form specific intra- and extracellular structures supporting cell structure when water is scarce (Richaud et al., 2020). Further, radical ions and reactive oxygen species can be sequestered by some IDPs, thereby mitigating oxidative stress. TDPs were initially characterized by their location of expression in the cell. Yamaguchi et al. (2012) used green fluorescent protein analysis to confirm the cytosolic abundant heat soluble (CAHS) TDP tended to gravitate toward the matrix of the cytoplasm. On the other hand, the secretory abundant heat soluble (SAHS) TDP was detected in the culture medium, indicating the protein had crossed the plasma membrane. As implied by the practice of naming TDPs simply for where they localize (Yamaguchi et al., 2012), there is, as yet, a paucity of research exploring the intricate mechanisms by which these proteins operate. This study, therefore, focuses on intrinsically disordered proteins, with an emphasis on tardigrade-specific cases.

### Study Overview

This study uses machine learning to uncover previously unidentified TDPs—as well as IDPs not exclusive to the phylum—to generate a novel list of high-priority proteins in need of further investigation. This research takes into consideration disorder metrics and phenotypic properties derived from literature curation and is novel in (1) comprehensively mining tardigrade proteomes for IDPs involved in cryptobiosis, (2) reporting, for the first time, a deduplicated and highly organized tardigrade pan-proteome database, and (3) crafting a concise list of proteins of interest in the anhydrobiotic phenotype. We anticipate that this nonredundant and centralized database, coupled with our novel machine learning pipeline, will serve as an asset for tardigrade researchers, and that the targeted list of proteins generated will grant future studies more direction.

## Project Goal

The overarching goal of this study was to strategically mine the tardigrade proteome, by way of machine learning, for intrinsically disordered proteins involved in the cryptobiotic phenotype.

## Objectives

1. To construct and deduplicate a comprehensive tardigrade pan-proteome database (PPD)
2. To use an intrinsic disorder prediction server (i.e., DISOPRED3) for analyzing location and intensity of disorder in proteins from the PPD
3. To employ a combination of machine learning and literature curation to filter disorder prediction results in order to yield a list of proteins in need of further exploration as prospective cryptobiosis-associated proteins
4. To utilize InterPro, a bioinformatic tool for functional analysis, to annotate the protein list generated

## Methodology

### Pan-Proteome Database Construction

#### Data Sourcing

In order to maximize the likelihood of detecting TDPs through disorder prediction and machine learning, tardigrade proteomic data were consolidated. First and foremost, a table was created encompassing all publicly available tardigrade protein sequences from UniProt (Universal Protein Resource (UniProt), 2021) and NCBI (National Center for Biotechnology Information (NCBI) Resource Coordinators, 2016). To eliminate redundancy, global pairwise sequence alignments were performed between NCBI and UniProt sequences using Harvard’s supercomputer cluster, O2. Only 104 NCBI sequences were able to be mapped (overlapped) as fragments of UniProt sequences, and 6,000+ unique NCBI protein sequences were added to the UniProt table. These merged datasets produced a table just under 45,000 proteins long. Experimental detection of proteins was noted in this supplementary table. Specifically, relevant peptide sequences were searched for in articles published between 2010 and 2017. Data from six articles were selected: Kamilari, Jørgensen, Schiøtt, and Møbjerg (2019), Schokraie et al. (2010), Schokraie et al. (2011), Schokraie et al. (2012), Yamaguchi et al. (2012), and Yoshida et al. (2017).

#### Redundancy Elimination (Deduplication)

Because these datasets were so vast (44,836 sequences), Excel was used to remove redundant sequence entries.

#### Sequence Mapping

To avoid introducing redundancy, to map between identical sequences cataloged under different names, and to maximize data inclusivity, literature-derived sequences were compared to sequences from UniProt and NCBI. A regular expression-based algorithm was devised in Perl: a brute force strategy that enumerates all possibilities for an edit distance (*ED*) between two sequences being aligned. For small *ED*, this algorithm runs faster than global sequence alignment and is therefore well-suited for mapping between nearly identical sequences for large datasets. Edit number dictated how sequences were dealt with (Figure 3).

**Figure 3.**
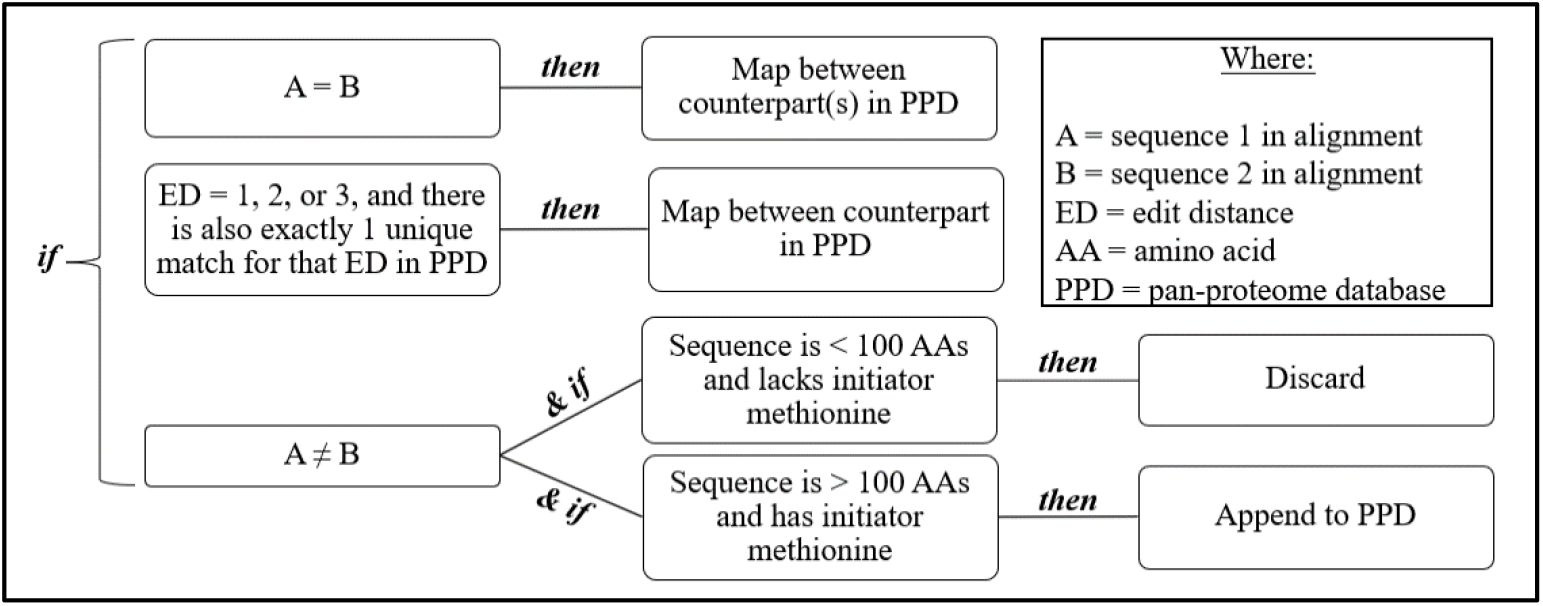
This decision tree illustrates how sequences were dealt with depending on the output of the regular expression script. Sequences were either mapped to the PPD, appended to the PPD as a unique addition, or discarded due to failure to meet minimum length requirements. Sequences with *ED* of 1-3 were included in the PPD because these small variations likely account for substitution mutations or minor mass spectrometry errors. Peptides < 100 AAs were discarded because they were not long enough for disorder prediction nor comparable to the training set. The initiator methionine presence requirement ensured proteins added to the PPD were not fragments.

### Intrinsic Disorder Prediction Utilizing DISOPRED3

DISOPRED3 is an intrinsic disorder prediction server that makes a binary call between ordered and disordered residues. DISOPRED2, its predecessor, was trained on sequences associated with missing residues in X-ray crystallography structures, a telltale sign of an intrinsically disordered region (IDR; Jones & Cozzetto, 2015). DISOPRED3 was selected because it was ranked highly in the 2014 Critical Assessments of Techniques for Protein Structure Prediction evaluation (Monastyrskyy, Kryshtafovych, Moult, Tramontano, & Fidelis, 2014). All PPD proteins were subjected to DISOPRED3 assessment.

### Machine Learning

Currently in the field of tardigrade research, there is no preemptive full-scale TDP identification system in place. Given the expensive and time-consuming nature of wet lab procedures for measuring protein expression and characterizing protein function, there is a chance TDPs that could be used to transfer tardigrade resilience to other species have yet to be identified, which compelled us to create the machine learning pipeline presented here as a means of automating the TDP identification process.

#### Feature Construction and IDP Metric Compilation

There is no single accepted set of guidelines for distinguishing IDPs from ordered proteins, ordered regions, or IDRs, so an additional literature search was conducted to locate common IDP characteristics. From this search, certain guiding principles were gleaned, namely that IDPs tend to (1) contain fewer hydrophobic amino acids (Oldfield & Dunker, 2014), which impedes hydrophobic collapse; (2) contain higher concentrations of disorder promoting amino acids such as proline and serine (Atkins, Boateng, Sorensen, & McGuffin, 2015); (3) contain higher concentrations of aromatic residues such as phenylalanine, tryptophan, and tyrosine (Oldfield & Dunker, 2014); and (4) that IDRs of more than 30 residues are considered long (Mohan, Uversky, & Radivojac, 2009). Based upon this literature search, the following dimensions were established for the disorder classifier: protein length, number of hydrophilic/hydrophobic residues, frequency of proline, glutamic acid, serine, glutamine, lysine, phenylalanine, tryptophan, tyrosine, and continuously disordered region length. Hydropathy determination was in accordance with Kyte-Doolittle (Kyte & Doolittle, 1982) and Hopp-Woods (Hopp & Woods, 1981) scales. Python was used to parse the long DISOPRED3 file and produce a summary output file recording the above-mentioned disorder statistics.

#### Training Sets

Negatives (ordered proteins) were flagged in the PPD based on if they had (1) a crystal structure in the Protein Data Bank (PDB; though this only accounted for three in the PDB, as accessed in October of 2020) and (2) enzyme commission (EC) numbers, as most enzymes have stable tertiary structures. Others were manually annotated, relying on confirmation that there were ordered PDB homologs. Positives (ordered proteins) were flagged in the PPD by searching the accompanying NCBI or UniProt annotation methodically for TDP keywords (i.e., CAHS, SAHS, MAHS, LEA, Dsup, and associated abbreviation expansions).

#### Support Vector Machine

Once training sets were established, it was necessary to select a machine learning model. When graphing a decision boundary, as the number of features, or dimensions, describing each data point increases (here, there were 11), the difficulty of separating data (44,836 points in the PPD) cleanly into distinct classes also rises. This “Curse of Dimensionality” (Bellman, 1966) is why we turn to artificial intelligence. This study utilized supervised machine learning in the form of a support vector machine (SVM). SVMs are highly effective classification tools with a computational edge over alternative separation techniques (Cristianini & Shawe-Taylor, 2000). The present SVM was created by modifying a preexisting code from Chang et al. (2020).

One challenge encountered while training the model was a severe class imbalance (55 entries in the positive training set versus 2,500+ entries in the negative training set). To account for this issue, 116 ordered examples were randomly selected to soften the imbalance while continuing to reflect size differences between the two sets. The balance issue was also dealt with using weighted scoring in the code itself. The scalar magnitude of the hyperplane equation’s coefficients for the resulting classifier (Table 1, overleaf) served as a measure of relative influence for each SVM feature. A linear kernel was utilized so coefficient values would be more readily interpretable. Also, feature values ranged drastically in magnitude. Therefore, standardization was performed through Z-scoring, mean centering the value distribution at zero and causing standard deviation of the transformed distribution to equal one. This aided hyperplane parameterization by preventing bias toward a certain scale. Gradient descent was employed to optimize hyperplane parameterization, running for between 100 and 1,000 iterations with the squared hinge loss function. This methodology combatted overfitting, given the traditional bias-variance tradeoff that plagues machine learning (Luxburg & Schölkopf, 2008).

**Table 1.** The hyperplane coefficient for each respective SVM feature. A higher coefficient magnitude indicates that its associated feature had a relatively higher influence on classification.

| Feature | Hyperplane Equation Coefficient |
| --- | --- |
| % hydrophobic residues | -0.65700478 |
| % proline | -0.42631545 |
| % hydrophilic residues | -0.38417876 |
| % glutamine | 0.35759344 |
| % lysine | 0.28354982 |
| % glutamic acid | 0.24527117 |
| % serine | -0.13962215 |
| % phenylalanine | -0.1332967 |
| % tyrosine | -0.10694951 |
| % disordered residues | -0.04497865 |
| % tryptophan | -0.04047342 |

Because the positive training set was derived from evolutionarily distinct groups (CAHS, SAHS, MAHS, LEA, and Dsup), they were not sufficiently homologous to each other to warrant using alignment scores as SVM features (Hesgrove & Boothby, 2020). To clarify, homology exists within each TDP class (e.g., CAHS versus CAHS), but less so across TDP classes (e.g., CAHS versus SAHS). Using alignment data in a classifier could have diluted its predictive power. Instead, the model was trained to generically predict TDPs. Proteins were subclassified afterward by performing ~300,000 local sequence alignments between positive predictions and positive training set proteins to classify predictions into their respective TDP subgroups. Bit scores for alignments between predicted positives and known positives were averaged with respect to each TDP subgroup, with the predicted subgroup being declared according to the highest average. Specifically, the BLOSUM62 standard substitution scoring matrix (Henikoff & Henikoff, 1992) was used, but with a more severe gap penalty, tailoring the code to locate loose evolutionary relationships (the penalty for a gap of any length was −5, and the added penalty for each residue greater than one for a gap was −1).

### Statistical Analysis

To assess how well the model separated the training set and generalized to unseen data, leave-one-out cross-validation was implemented. Afterward, to ensure cross-validation results weren’t rendered by chance, additional leave-one-out cross-validation rounds were conducted (cross-validation confirmation), first with a shuffled feature space, and then again with shuffled training set class labels (a randomized control). Subclass prediction confidence was also evaluated. 55 known TDPs from the positive training set were aligned against each other. Then, bit scores from within each class (e.g., CAHS versus CAHS) and across classes (e.g., CAHS versus SAHS) were recorded in Excel to locate a class-specific threshold separating bimodal distributions (Table 2, overleaf). To be deemed a confident prediction, the predicted positive bit score had to meet the threshold for its respective predicted class. Though this procedure does not produce an e-value or p-value, it does indicate the confidence of each predicted positive, erring on the more conservative side, as it corresponds to a false positive rate of 0%, with respect to TDP subtypes.

**Table 2.** The minimum local alignment bit score required for a protein to be deemed confidently a member of its predicted subclass. Bit score average with respect to each subclass for each disordered protein prediction was compared to the threshold for its associated subclass. If the bit score average was greater than the threshold, it was deemed a confident subclass prediction.

| Subclass | CAHS | SAHS | MAHS | LEA | Dsup |
| --- | --- | --- | --- | --- | --- |
| Threshold | 502 | 129 | 146 | 502 | 329 |

## Results

### SVM Performance

The classifier designed in this study was proficient at separating known TDPs from known non-TDPs in cross-validated training set performance, correctly classifying 160/171 training set proteins, yielding raw accuracy of ~93.6% (Figure 4). This points to the suitability of disorder and sequence features selected for SVM training. For training set performance, the area under the curve (AUC) score for the receiver operating characteristic (ROC) curve was ~0.98. To confirm the hyperplane was not overfitted to training set data, leave-one-out cross-validation was performed. The leave-one-out cross-validated AUC of the ROC was ~0.95. To ensure the promising cross-validation results were not generated by chance, secondary cross-validation (confirmation) was conducted, which involved rerunning cross-validation, but with shuffled training data. For the first round of cross-validation confirmation, with training on shuffled labels, the AUC score was ~0.56. For the second round, with training on shuffled feature space, the AUC score was ~0.52 (Figure 5). Shuffled training data diminished the predictive capacity of the SVM, as expected. Qualitatively speaking, the closeness of model performance during training and leave-one-out cross-validation alludes to negligible overfitting and the strong, generalizable predictive power of the overall pipeline.

**Figure 4.**
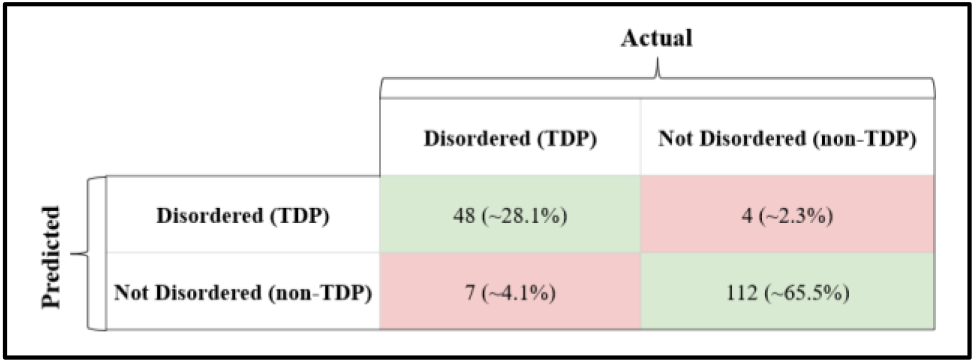
Confusion matrix illustrating (counterclockwise from upper left) the proportion of true positives, false negatives, true negatives, and false positives for the SVM’s training set of 171 proteins.

**Figure 5.**
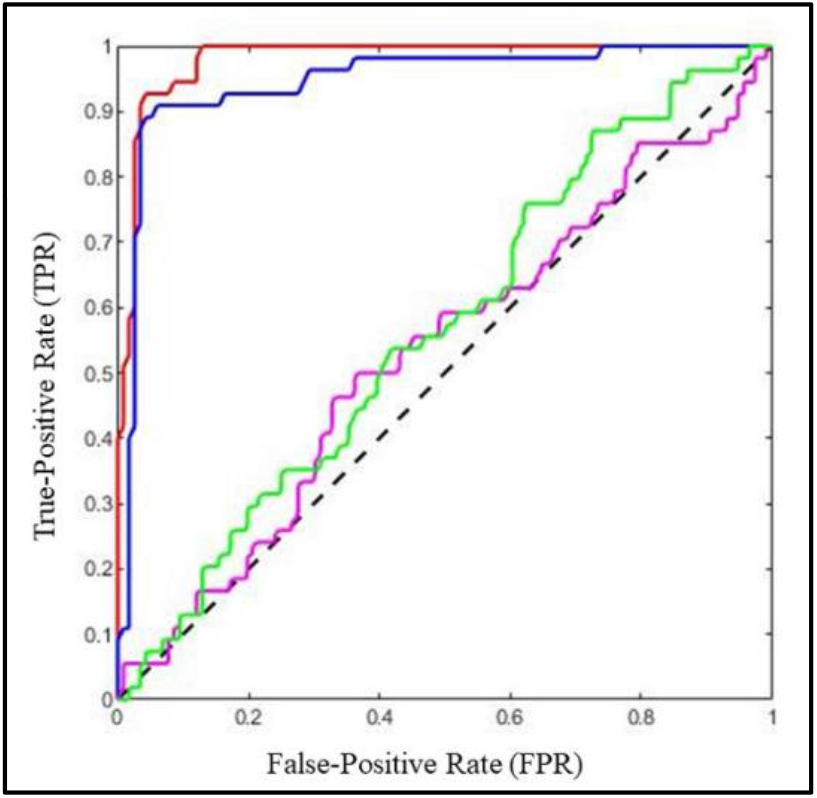
Red: SVM training set ROC (AUC ≈ 0.98). Blue: leave-one-out cross-validated ROC (AUC ≈ 0.95). Magenta: shuffled features ROC (AUC ≈ 0.52). Green: shuffled labels ROC (AUC ≈ 0.56).

### Finalized, High-Priority List

At the core of this study was the creation of a concise list of TDPs intended to guide researchers in the budding field of tardigrade-based technology development. The SVM predicted 5,415 previously unknown IDPs/TDPs out of 44,836 total PPD proteins. Of these 5,415 proteins, 52 were also detected experimentally in previous studies, meaning they map to sequences from the six publications listed under “Data Sourcing.” Also of these 5,415 proteins, 30 passed the bit score threshold for homology to TDP subclasses. To gain insight into how these proteins might play a role in cryptobiosis, InterPro, a protein annotation tool (Apweiler et al., 2001; Blum et al., 2021), was used to analyze the 82 proteins (Appendix). Many entries were “hypothetical proteins,” which we were able to characterize with InterPro, shedding light on their function. This narrowed down list will empower researchers to sidestep tedious experimental sifting for disordered proteins.

### InterPro Annotations

Proteins in the finalized list were manually annotated with InterPro to provide a foothold for scientists exploring TDPs, as well as to expedite the genesis of new TDP technology. InterPro annotations showed that the 82 proteins of interest are involved in diverse biological processes, including motor activity, ATP binding, DNA binding, and ion sequestration. Curiously, 15/82 (~18.3%) were annotated as heat shock proteins (Hsp20, Hsp40, Hsp70, and Hsp90 were all detected). In this group, prevalence of heat shock proteins, a type of molecular chaperone that assists in biomolecule assembly/disassembly, reinforces their role in the tardigrade cryptobiotic molecular landscape (Schokraie et al., 2010; Schokraie et al., 2011). Characterizing these putative TDPs is a pivotal step forward in this field.

### Subclassification Trends

For TDP subclassification, 297,825 local sequence alignments were performed (55 training set proteins x 5,415 positive predictions). The 52 predictions with experimental detection in previous literature were not deemed confident by way of threshold comparison, though it should be noted that the threshold confidence metric errs on the conservative side (corresponding to a false positive rate of 0%). Of the 5,415 disordered predictions, 30 subclass assignment bit score averages met the threshold. This confidence measurement does *not* apply to the disordered prediction of the protein, but exclusively to subclass assignment (e.g., CAHS versus SAHS). Most positives were subclassified as damage suppressing proteins (Figure 6). The bit score distributions utilized to determine the class-specific thresholds are displayed in Figure 7 (overleaf).

**Figure 6.**
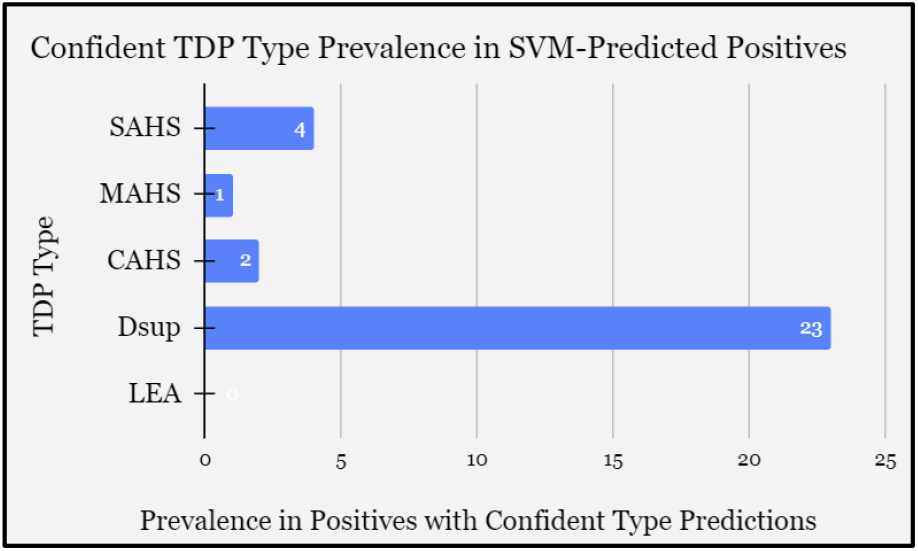
Based on bit score averages from local alignments between positive predictions and the positive training set, these bar graphs depict how many positive predictions were assigned to each subclass (SAHS, MAHS, CAHS, Dsup, and LEA) of the 30 positives with confident subclass predictions.

**Figure 7.**
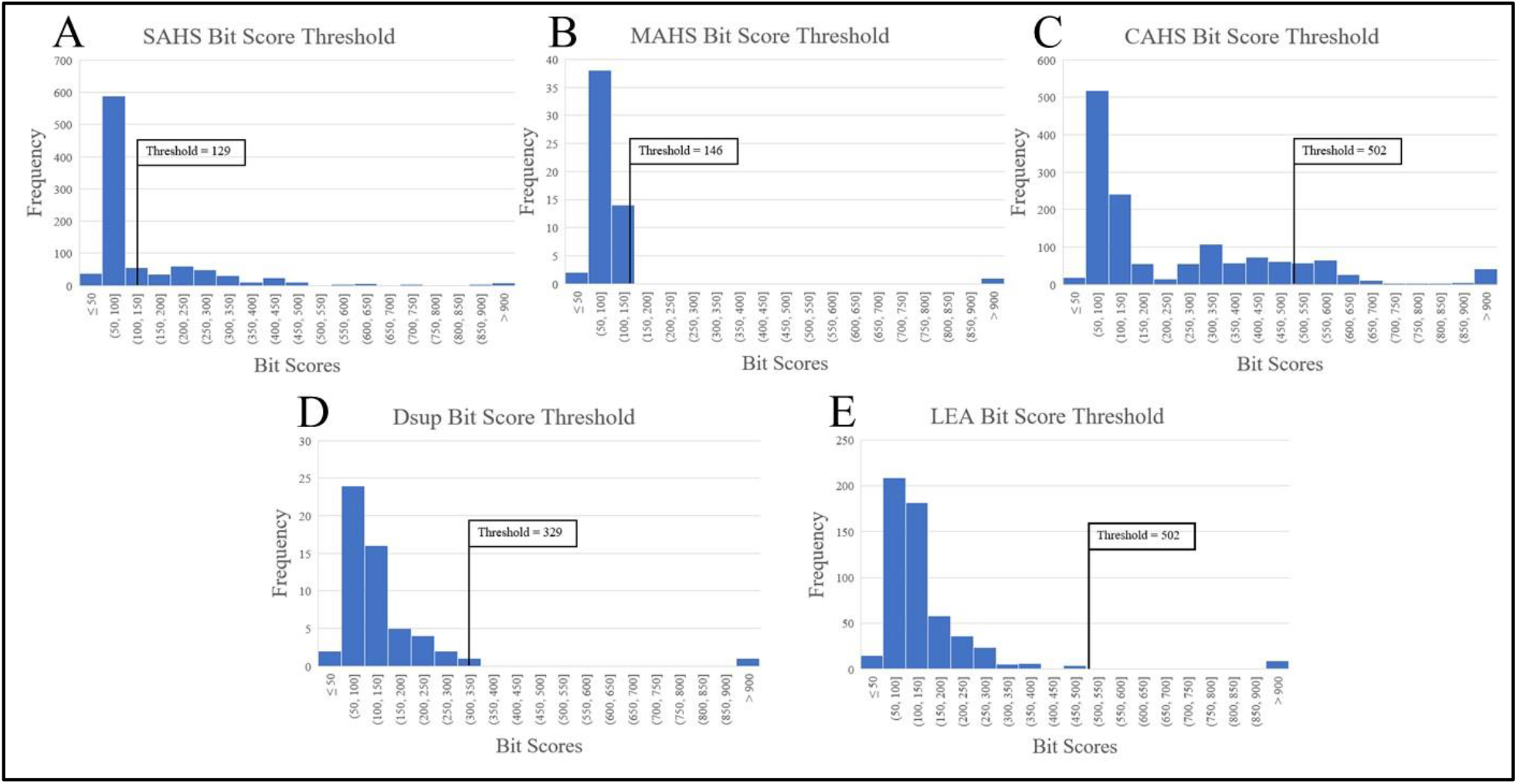
Graphs A-E show bit score distributions analyzed to determine class-specific confidence thresholds. After predicted and known positives were subject to local sequence alignments, the resultant bit scores were averaged with respect to each subclass, and the subclass associated with the highest average for each protein was deemed its predicted subclass. The confidence threshold represents the highest bit score that can be achieved from an alignment between two sequences from unrelated subclasses. As demonstrated by how the majority of bit scores lie below the threshold, this confidence metric is quite rigorous.

## Discussion

### A Tool for More Targeted Research

One immediate application of this classifier is compensating for both the deficit in and drawbacks of tardigrade experiments. The handful of existing publications we pulled data from, though carefully selected, presumably reflect a fraction of tardigrade protein expression under various stress conditions. Moreover, IDPs are difficult to measure and identify because many can form higher-order structures that can may make them inaccessible to standard platforms such as nuclear magnetic resonance (Radivojac et al., 2004) or x-ray crystallography. Now that we have narrowed down proteins of interest, these can be targeted for specific measurements experimentally, such as with mass spectrometry, or genetic experiments. Since it would be a costly and lengthy process to investigate all PPD proteins experimentally, identifying proteins of interest beforehand ensures resources are spent prudently. While predictive pipelines are typically utilized to weed out IDPs and focus on ordered proteins (Atkins et al., 2015), the same principle can be readily reversed, as was done here, for the purpose of studying IDPs.

### Making Sense of Disorder

IDPs are vital to extremotolerance and cell biology in general (Dunker, Bondos, Huang, & Oldfield, 2015; Wright & Dyson, 2015), playing crucial roles in protein-protein interactions due to their flexibility, but they are severely understudied (Necci, Piovesan, Critical Assessment of Protein Intrinsic Disorder (CAID) Predictors, Database of Protein Disorder (DisProt) Curators, & Tosatto, 2021).

Identifying illustrative examples of IDPs here paves the way for more deeply understanding them. This study expands the number of known examples and thereby potentiates further analysis into what constitutes an IDP. The 5,415 SVM positive predictions, if they are indeed disordered, dramatically enlarge the register of known tardigrade-specific IDPs (TDPs), which is a notable step forward for creating TDP technologies with impacts in fields ranging from astrobiology to genetic engineering.

### Next Steps

An area for additional investigation would be constructing a separate SVM classifier for each TDP subclass. Future studies could also increase the number of shuffled cross-validation confirmation trials and develop a system for strategically, instead of randomly, selecting the negative training set. To fully reap the benefits of TDP-based technology, researchers need to assess proteins we identified *in vitro*, such as through mass spectrometry and western blotting (as in Schokraie et al., 2010), as determining which of these potential IDPs are upregulated during desiccation, and to what extent, will be critical for identifying with confidence which are ideal for incorporation into biotechnology. Other avenues of investigation would include designing genetic knockout/knockdown experiments in tardigrades for targeting proteins identified here, as well as testing for transfer of stress tolerance to heterologous systems and performing stress tolerance assays.

### Tardigrade-Specific Intrinsically Disordered Proteins and Biotechnology

Recent studies have observed how TDPs can be utilized as a means for transferring oxidative stress tolerance (Chavez et al., 2019), radiation tolerance (Hashimoto et al., 2016; Kirke, Jin, & Zhang, 2020; Westover et al., 2020), and osmotic stress tolerance (Tanaka et al., 2015) between organisms. Notably, in 2017, transfer of CAHS proteins to yeast increased their desiccation tolerance approximately 100-fold (Boothby et al., 2017). The same study found *Escherichia coli* exhibited a similar response, pointing to the promise of using TDPs to confer tardigrade extremotolerance to other life forms.

Therefore, while this study fortifies knowledge of basic tardigrade biology and stress response, it has ramifications for higher life forms as well. Serendipitously, certain proteins pinpointed were likened to those found in humans. Common examples from the Appendix include fatty acid-binding proteins (Smathers & Petersen, 2011) and DnaJ/DnaK proteins. Table 3 presents select proteins from the Appendix and their human homologs, logical candidates for technological applications.

**Table 3.**
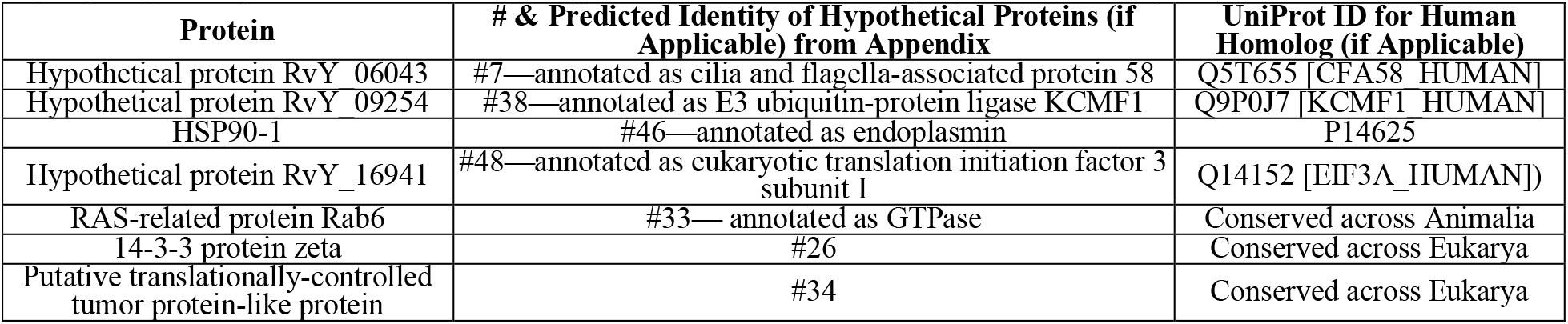
Highlighting SVM-predicted TDPs from the Appendix with human homologs (when applicable).

Insofar as TDPs take on a material state upon desiccation (Boothby et al., 2017), form supportive cell structures (Richaud et al., 2020), and sequester ions (Bray, 1993), they hold great promise for many advantageous applications. To clarify, TDPs could be used to design more effective, affordable biomaterial preservation methods. The concept of mimicking anhydrobiotic creatures to preserve organs, tissues, and blood has been pondered for half a century (Keilin, 1959). Today, cytosolic abundant heat-soluble TDPs (CAHS) are already being evaluated for manufacturing pharmaceutical excipients, such as lactate dehydrogenase and lipoprotein lipase, and the results are encouraging (Piszkiewicz et al., 2019). Such a preservative could also aid storage of medical supplies containing biological components, such as vaccines. To that end, because CAHS proteins protect enzymes from desiccation and lyophilization, they have been tested as a vaccine stabilizer. When a vaccine with this preservative was tested in mice, the stabilizer was deemed as not toxic, and treatment elicited antibody production as intended (Esterly et al., 2020). Cold storage is sometimes a limiting factor when it comes to distributing vaccines (Kaufmann, Miller, & Cheyne, 2011; Zaffran et al., 2013)—such as with some COVID-19, chicken pox, and Ebola vaccines—but with TDP stabilizers, certain vaccine stockpiles could be dispersed and stored safely and affordably, circumventing the cumbersome cold chain and even ultra-cold chain vaccine preservation methods, thus increasing global medical equity.

Outside of the realm of biopreservation, TDPs could be a valuable addition to therapeutics. For instance, they could be incorporated into treatments for delaying deleterious cell processes when an injury occurs in an isolated location. Delaying necrosis and apoptosis would be especially germane to improving patient health outcomes when treatment is time-sensitive. This advancement could be crucial for the military in decreasing battlefield mortality.

TDPs could also have applications in medical conditions involving oxidative stress. Because oxidative stress can contribute to cancer (Liguori et al., 2018), antioxidant properties of these proteins (Hashimoto et al., 2016; Rizzo et al., 2010) could prove useful as part of new treatments. Specifically, certain tardigrade proteins form structures that can capture reactive oxygen species and cut short their damaging ripple effect within the cell.

Aside from medical implications, mimicking tardigrade resilience has more broad technological applications, namely in coping with Earth’s changing climate. For instance, a damage suppressor protein (Dsup) gene was expressed in plants and decreased radiation-induced DNA damage in Kirke et al. (2020). Theoretically, extremotolerant plants could be engineered to maintain a stable food supply despite escalating environmental challenges. In the far future, perhaps such recombinant plants could play a role in sustaining life on other planets.

On the note of space travel, tardigrades have survived the brutal conditions of space on multiple missions since the early 2000s (Jönsson et al., 2008; Persson et al., 2011; Rebecchi et al., 2009, 2010, 2011; Vukich et al., 2012) and are considered model organisms for space research (Bertolani et al., 2001; Guidetti, Rizzo, Altiero, & Rebecchi, 2012; Jönsson, 2007; May, Maria, & Guimard, 1964). Seeing as certain TDPs are involved in radiotolerance (Hashimoto et al., 2016), they could be used to engineer radiation-resistant crops or even to protect astronauts against cosmic radiation, helping them explore new frontiers.

## Conclusion

Researchers have long marveled at tardigrade resilience, instigating today’s endeavors in tardigrade biomimicry (the imitation of biological phenomena in man-made technology). Given mounting evidence that intrinsically disordered proteins (IDPs) play a pivotal role in cryptobiotic phenomena, the dearth of literature on IDPs is peculiar. In this study, a computational method for identifying tardigrade-specific IDPs (TDPs) of interest was devised. This paves the way for potentially revolutionary TDP/IDP-based technology. The novel and thorough pan-proteome database (PPD) presented here yielded a prioritized shortlist of hypothetically cryptobiosis-associated IDPs, which were identified by a novel support vector machine classifier trained on IDP properties. Our classifier performed promisingly well, accurately classifying ~93.6% of the training set. During testing, 5,415 of the 44,836 PPD proteins were categorized as disordered. InterPro annotations of 82 positives, some heretofore unreported in the literature, suggest a plausible facilitative role for many of them in tardigrade extremotolerance. By methodically funneling the expansive tardigrade proteome into a more succinct proteins of interest list, this study charts a course for more directed research into tardigrade cryptobiosis, potentially permitting humans to one day exploit and cross-apply the tardigrade’s natural strengths for the benefit of such fields as medicine, biotechnology, and even space travel. Further, considering how water shortages have wreaked havoc on crops and livestock in the wake of climate change, the prospect of engineering TDP-expressing crops that can withstand drought is of special import. The tardigrade’s perplexing proteins could be key to conferring the same desiccation tolerance that has been observed in the enigmatic tardigrade for hundreds of years. Ironically, unraveling the underpinnings of protein disorder could be paramount for creating a more ordered world.

## Supporting information

Tardigrade Pan-Proteome Database

## Acknowledgements

High-throughput computation was enabled by the Harvard Medical School O2 high performance compute cluster.

## Appendix

### Catalog of 82 Putative Intrinsically Disordered Proteins

Greater distance to hyperplane indicates more confident classification. Predicted Type column is based on bit score averages from local alignments between predicted/known positives. Amino acid is abbreviated as AA throughout. 52 red entries: experimentally detected in previous studies, with predicted types not satisfying bit score significance threshold. 30 blue entries: confident TDP type predictions without experimental detection.

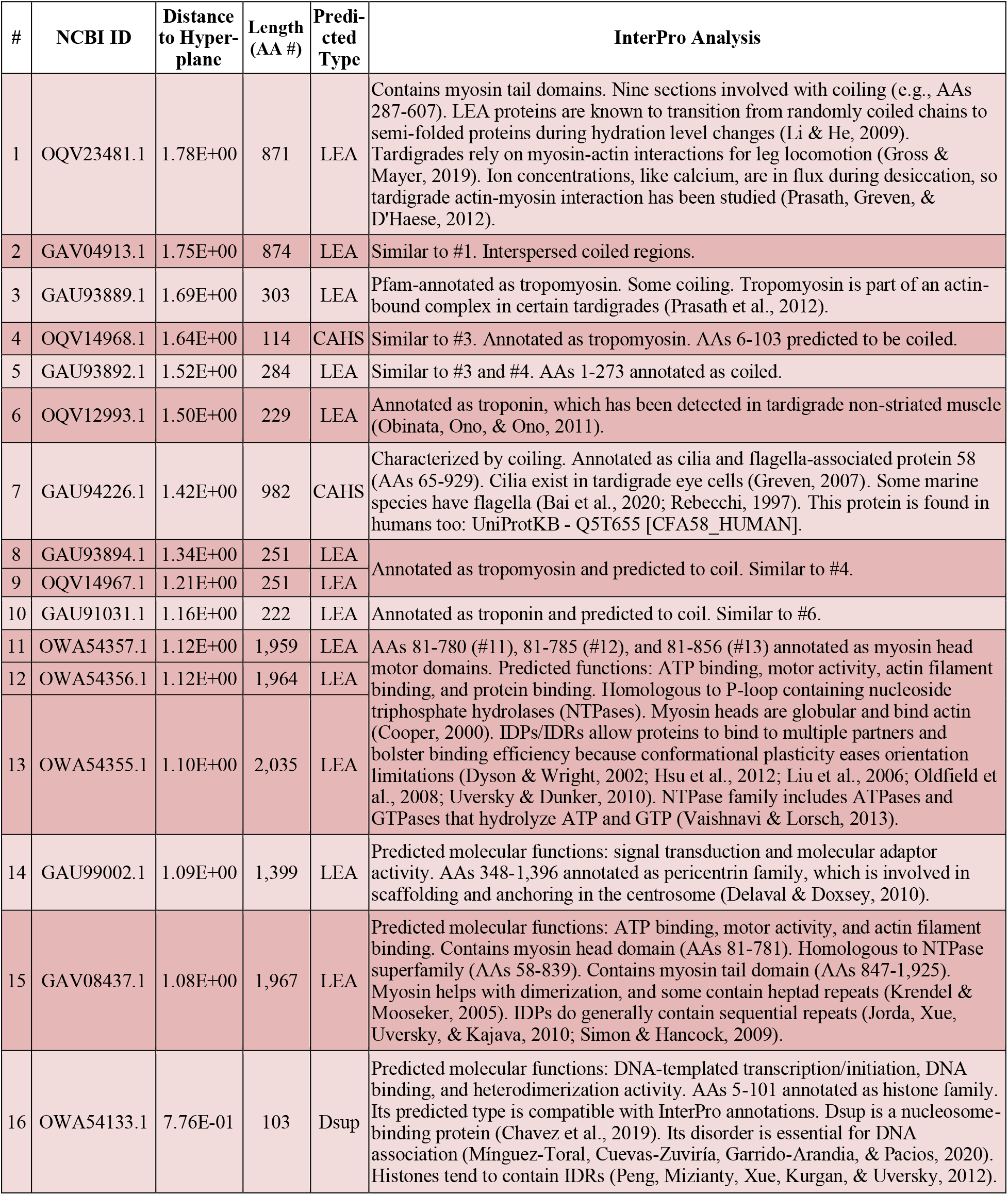
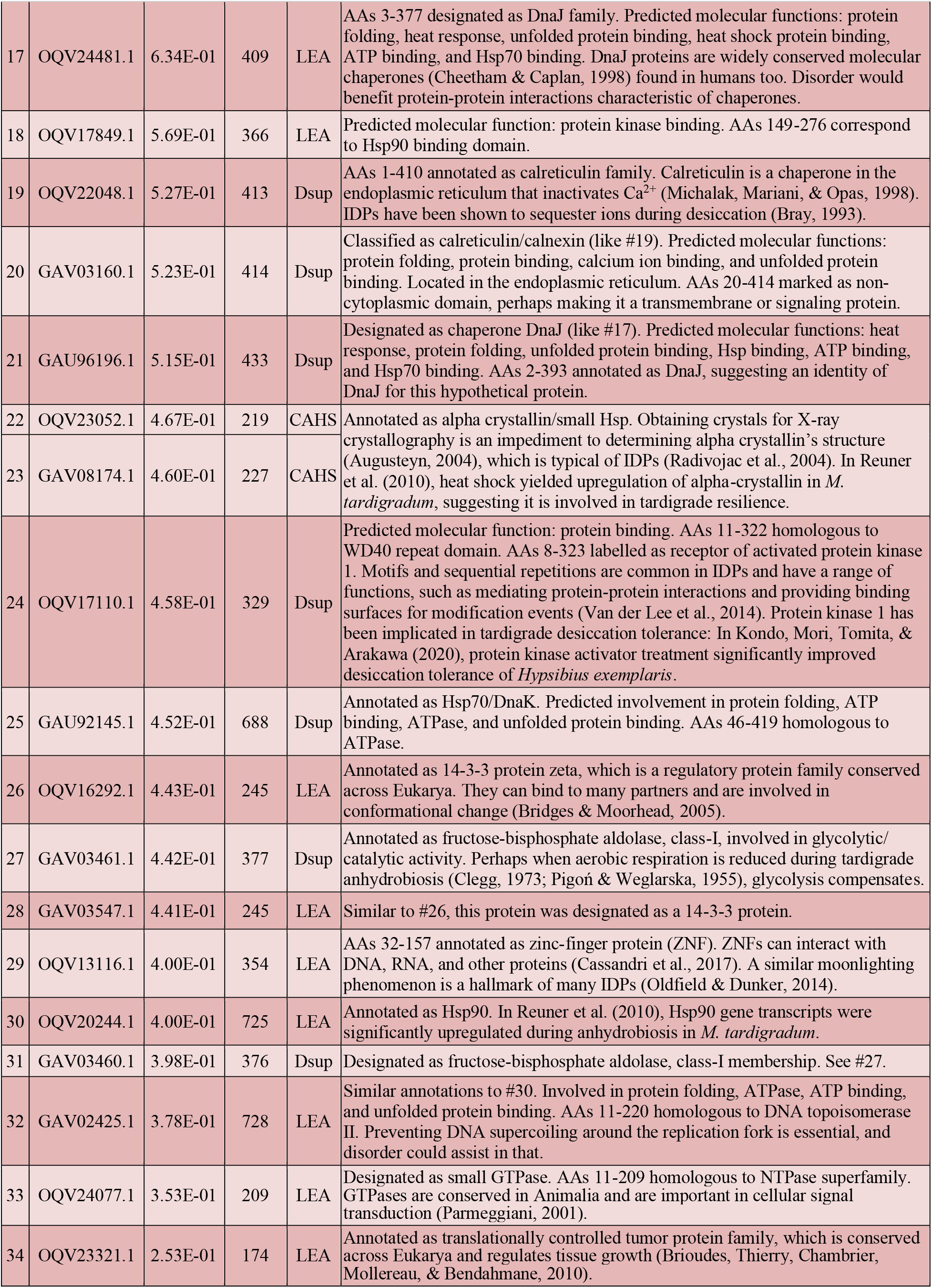
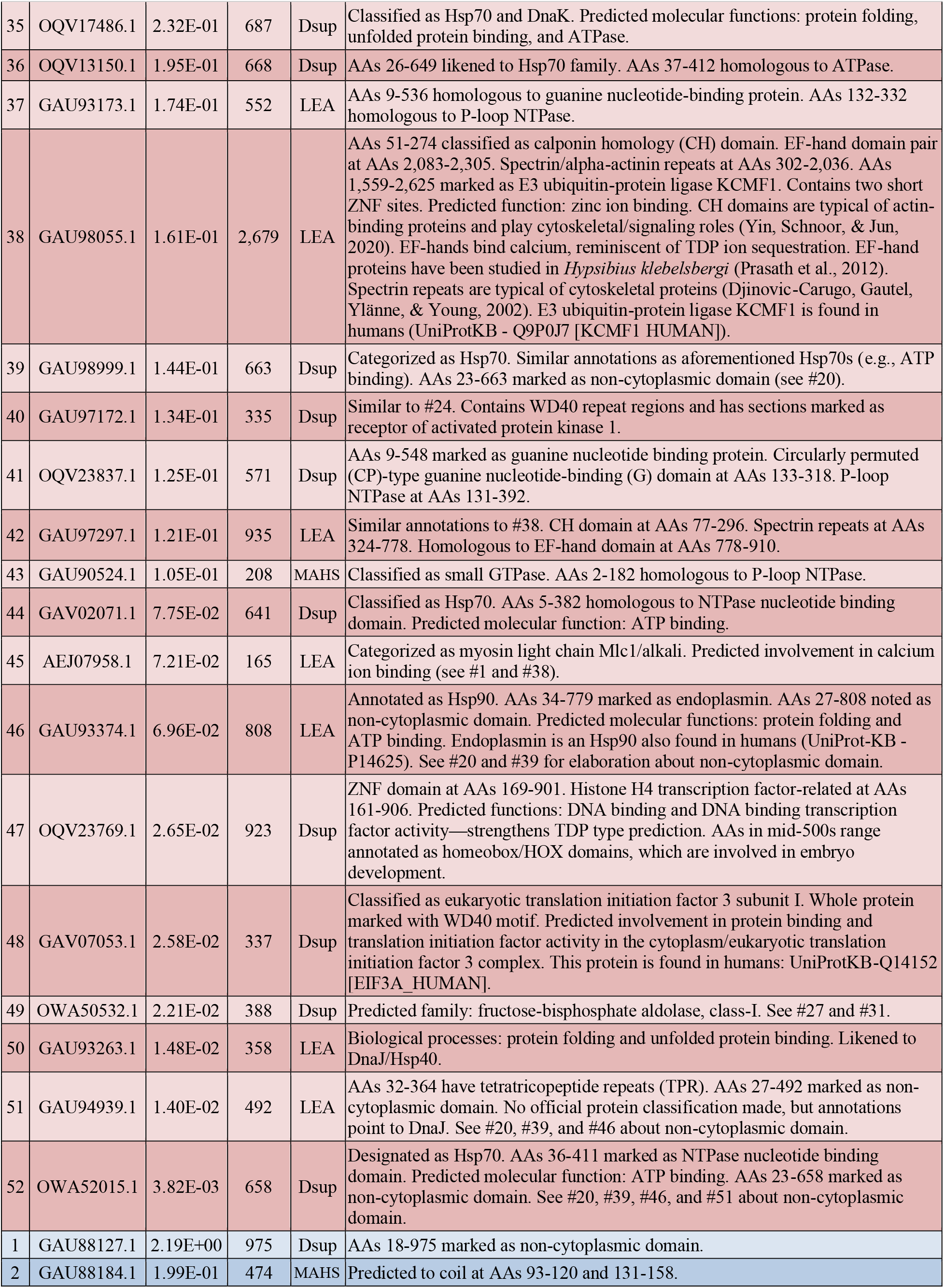
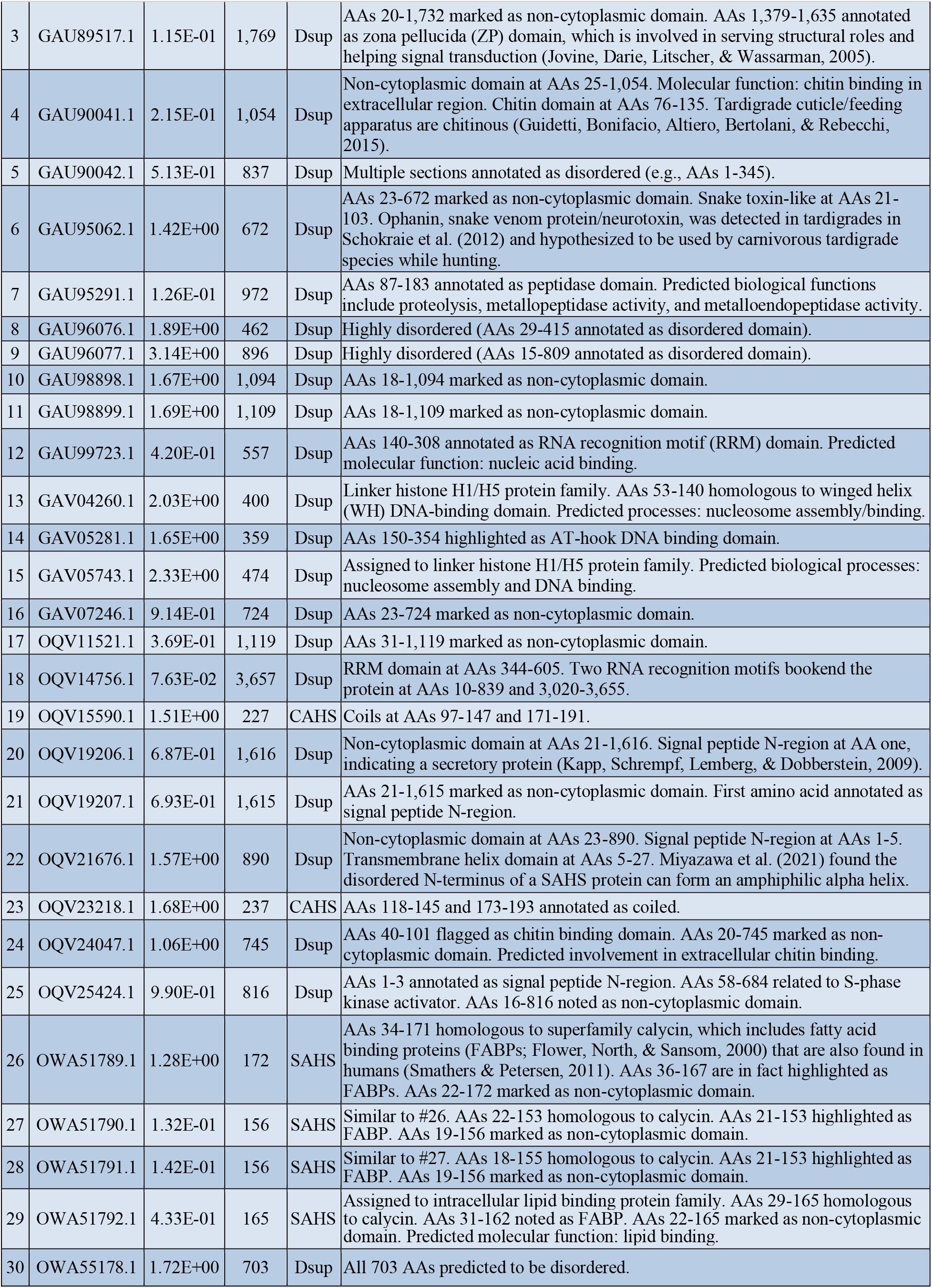

## Notes

### Competing Interest Statement

The authors have declared no competing interest.

## References

Alterio, T., Guidetti, R., Boschini, D., & Rebecchi, L. (2012). Heat shock proteins in encysted and anhydrobiotic eutardigrades. Journal of Limnology, 71(1), e22. https://doi.org/10.4081/jlimnol.2012.e22

Apweiler, R., Attwood, T. K., Bairoch, A., Bateman, A., Birney, E., Biswas, M., Bucher, P., Cerutti, L., Corpet, F., Croning, M. D., Durbin, R., Falquet, L., Fleischmann, W., Gouzy, J., Hermjakob, H., Hulo, N., Jonassen, I., Kahn, D., Kanapin, A., Karavidopoulou, Y., Lopez, R., Marx, B., Mulder, M. J., Oinn, T. M., Pagni, M., Servant, F., Sigrist, C. J. A., & Zdobnov, E. M. (2001). The InterPro database, an integrated documentation resource for protein families, domains and functional sites. Nucleic Acids Research, 29(1), 37–40. https://doi.org/10.1093/nar/29.1.37

Atkins, J. D., Boateng, S. Y., Sorensen, T., & McGuffin, L. J. (2015). Disorder prediction methods, their applicability to different protein targets and their usefulness for guiding experimental studies. International Journal of Molecular Sciences, 16(8), 19040–19054. https://doi.org/10.3390/ijms160819040

Augusteyn, R. C. (2004). Alpha-crystallin: A review of its structure and function. Clinical and Experimental Optometry, 87(6), 356–66. https://doi.org/10.1111/j.1444-0938.2004.tb03095.x

Bai, L., Wang, X., Zhou, Y., Lin, S., Meng, F., & Fontoura, P. (2020). *Moebjergarctus clarionclippertonensis*, a new abyssal tardigrade (Arthrotardigrada, Halechiniscidae, Euclavarctinae) from the Clarion-Clipperton Fracture Zone, North-East Pacific. Zootaxa, 4755(3). https://doi.org/10.11646/zootaxa.4755.3.8

Baumann, H. (1922). Die anabiose der tardigraden. Zoologische Jahrbücher, 45, 501–556.

Bellman, R. (1966). Dynamic programming. Science, 153(3731), 34–37.

Beltrán-Pardo, E., Jönsson, K. I., Harms-Ringdahl, M., Haghdoost, S., & Wojcik, A. (2015). Tolerance to gamma radiation in the tardigrade *Hypsibius dujardini* from embryo to adult correlate inversely with cellular proliferation. PLoS One, 10(7), e0133658. https://doi.org/10.1371/journal.pone.0133658

Bertolani, R., Rebecchi, L., Jönsson, K. I., Borsari, S., Guidetti, R., Altiero, T. (2001). Tardigrades as a model for experiences of animal survival in space. In R. Monti & C. Bonifazi (Eds.), La Scienza e la Tecnologia Spaziale sulla Stazione Internazionale (ISS) (2nd ed., pp. 211–212). Spec. Issue ASI Natl. Workshop, Turin, 2001. MSSU—Micro Space Station Util.

Blum, M., Chang, H. Y., Chuguransky, S., Grego, T., Kandasaamy, S., Mitchell, A., Nuka, G., Paysan-Lafosse, T., Qureshi, M., Raj, S., Richardson, L., Salazar, G. A., Williams, L., Bork, P., Bridge, A., Gough, J., Haft, D. H., Letunic, I., Marchler-Bauer, A., Mi, H., Natale, D. A., Necci, M., Orengo, C. A., Pandurangan, A. P., Rivoire, C., Sigrist, C. J. A., Sillitoe, I., Thanki, N., Thomas, P. D., Tosatto, S. C. E., Wu, C. H., Bateman, A., & Finn, R. D. (2021). The InterPro protein families and domains database: 20 years on. Nucleic Acids Research, 49(D1), D344–D354. https://doi.org/ https://doi.org/10.1093/nar/gkaa977

Bonnet, C. & Goeze, J. A. E. (1773). Herrn Karl Bonnets Abhandlungen aus der Insektologie. Halle: Bey J.J. Gebauers Wittwe und Joh. Jac. Gebauer. http://www.biodiversitylibrary.org/item/102801#page/7/mode/1up

Boothby, T. C., Tapia, H., Brozena, A. H., Piszkiewicz, S., Smith, A. E., Giovannini, I., Rebecchi, L., Pielak, G. J., Koshland, D., & Goldstein, B. (2017). Tardigrades use intrinsically disordered proteins to survive desiccation. Molecular Cell, 65(6), 975–984. https://doi.org/10.1016/j.molcel.2017.02.018

Bray, E. A. (1993). Molecular responses to water deficit. Plant Physiology, 103(4), 1035–1040. https://doi.org/10.1104/pp.103.4.1035

Bridges, D. & Moorhead, G. B. (2005). 14-3-3 proteins: A number of functions for a numbered protein. Science Signalling, 296, re10. https://doi.org/10.1126/stke.2962005re10

Brioudes, F., Thierry, A., Chambrier, P., Mollereau, B., & Bendahmane, M. (2010). Translationally controlled tumor protein is a conserved mitotic growth integrator in animals and plants. Proceedings of the National Academy of Sciences, 107(37), 16384–16389. https://doi.org/10.1073/pnas.1007926107

Cassandri, M., Smirnov, A., Novelli, F., Pitolli, C., Agostini, M., Malewicz, M., Melino, G., & Raschellà, G. (2017). Zinc-finger proteins in health and disease. Cell Death Discovery, 3, 17071. https://doi.org/10.1038/cddiscovery.2017.71

Chang, R. L., Stanley, J. A., Robinson, M. C., Sher, J. W., Li, Z., Chan, Y. A., Omdahl, A. R., Wattiez, R., Godzik, A., & Matallana-Surget, S. (2020). Protein structure, amino acid composition and sequence determine proteome vulnerability to oxidation-induced damage. EMBO Journal, e104523. https://doi.org/10.15252/embj.2020104523

Chavez, C., Cruz-Becerra, G., Fei, J., Kassavetis, G. A., & Kadonaga, J. T. (2019). The tardigrade damage suppressor protein binds to nucleosomes and protects DNA from hydroxyl radicals. eLife, 8, e47682. https://doi.org/10.7554/eLife.47682

Cheetham, M. E. & Caplan, A. J. (1998). Structure, function and evolution of DnaJ: Conservation and adaptation of chaperone function. Cell Stress Chaperones, 3, 28–36.

Clegg, J. S. (1973). Do dried cryptobiotes have a metabolism? In: Crowe JH, Clegg JS, eds. Anhydrobiosis. Stroudsburg: Dowden, Hutchinson and Ross, 141–147.

Cooper, G. M. (2000). The cell: A molecular approach (2nd ed.). Sunderland (MA): Sinauer Associates. https://www.ncbi.nlm.nih.gov/books/NBK9961/

Cristianini, N. & Shawe-Taylor, J. (2000). Support vector machines. In An Introduction to Support Vector Machines and Other Kernel-based Learning Methods (pp. 93–124). Cambridge: Cambridge University Press. https://doi.org/10.1017/CBO9780511801389.008

Crowe, J. H., Carpenter, J. F., & Crowe, L. M. (1998). The role of vitrification in anhydrobiosis. Annual Review of Physiology, 60, 73–103. https://doi.org/10.1146/annurev.physiol.60.1.73

Crowe, L. M. (2002). Lessons from nature: The role of sugars in anhydrobiosis. Comparative Biochemistry and Physiology - Part A: Molecular & Integrative Physiology, 131(3), 505–513. https://doi.org/10.1016/S1095-6433(01)00503-7

Degma, P., Bertolani, R., & Guidetti, R. (2021). Actual checklist of Tardigrada species. http://dx.doi.org/10.25431%2F11380_1178608

Delaval, B. & Doxsey, S. J. (2010). Pericentrin in cellular function and disease. The Journal of Cell Biology, 188(2), 181–190. https://doi.org/10.1083/jcb.200908114

Djinovic-Carugo, K., Gautel, M., Ylänne, J., & Young, P. (2002). The spectrin repeat: A structural platform for cytoskeletal protein assemblies. FEBS Letters, 513, https://doi.org/10.1016/S0014-5793(01)03304-X

Doyère, P. L. N. (1842). Memoires sur les tardigrade. Sur le facilité possedent les tardigrades, les rotifers, les anguillules de toit et quelques autre animalcules, de revenir à la vie après été completement déssechées. Annales des Sciences Naturelles. Zoologie et Biologie Animale, 18, 5–35.

Dunker, A. K., Bondos, S. E., Huang, F., & Oldfield, C. J. (2015). Intrinsically disordered proteins and multicellular organisms. Seminars in Cell & Developmental Biology, 37, 44–55. https://doi.org/10.1016/j.semcdb.2014.09.025

Dyson, H. J. & Wright, P. E. (2002). Coupling of folding and binding for unstructured proteins. Current Opinion in Structural Biology, 12, 54–60. https://doi.org/10.1016/S0959-440X(02)00289-0

Esterly, H. J., Crilly, C. J., Piszkiewicz, S., Shovlin, D. J., Pielak, G. J., & Christian, B. E. (2020). Toxicity and immunogenicity of a tardigrade cytosolic abundant heat soluble protein in mice. Frontiers in Pharmacology, 11, 565969. https://doi.org/10.3389/fphar.2020.565969

Flower, D. R., North, A. C., & Sansom, C. E. (2000). The lipocalin protein family: Structural and sequence overview. Biochimica et Biophysica Acta, 1482(1-2), 9–24. https://doi.org/10.1016/S0167-4838(00)00148-5

Förster, F., Beisser, D., Grohme, M. A., Liang, C., Mali, B., Siegl, A. M., Engelmann, J. C., Shkumatov, A. V., Schokraie, E., Müller, T., Schnölzer, M., Schill, R. O., Frohme, M., & Dandekar, T. (2012). Transcriptome analysis in tardigrade species reveals specific molecular pathways for stress adaptations. Bioinformatics and Biology Insights, 6, 69–96. https://doi.org/10.4137/BBI.S9150

Förster, F., Liang, C., Shkumatov, A., Beisser, D., Engelmann, J. C., Schnölzer, M., Frohme, M., Müller, T., Schill, R. O., & Dandekar, T. (2009). Tardigrade workbench: Comparing stress-related proteins, sequence-similar and functional protein clusters as well as RNA elements in tardigrades. BMC Genomics, 10, 469. https://doi.org/10.1186/1471-2164-10-469

Franceschi, T. (1948). Anabiosi nei tardigradi. Boll. Mus. Ist. Biol. Univ. Genova. 22: 47–49.

Greven, H. (2007). Comments on the eyes of tardigrades. Arthropod Structure & Development, 36(4), 401–407. https://doi.org/10.1016/j.asd.2007.06.003

Gross, V. & Mayer, G. (2019). Cellular morphology of leg musculature in the water bear *Hypsibius exemplaris* (Tardigrada) unravels serial homologies. Royal Society Open Science, 6(10), 191159. https://doi.org/10.1098/rsos.191159

Guidetti, R., Bonifacio, A., Altiero, T., Bertolani, R., & Rebecchi, L. (2015). Distribution of calcium and chitin in the tardigrade feeding apparatus in relation to its function and morphology. Integrative and Comparative Biology, 55(2), 241–52. https://doi.org/10.1093/icb/icv008

Guidetti, R., Rizzo, A. M., Altiero, T., & Rebecchi, L. (2012) What can we learn from the toughest animals of the Earth? Water bears (tardigrades) as multicellular model organisms in order to perform scientific preparations for lunar exploration. Planetary and Space Science, 74(1), 97–102. https://doi.org/10.1016/j.pss.2012.05.021

Hashimoto, T., Horikawa, D. D., Saito, Y., Kuwahara, H., Kozuka-Hata, H., Shin-I, T., Minakuchi, Y., Ohishi, K., Motoyama, A., Aizu, T., Enomoto, A., Kondo, K., Tanaka, S., Hara, Y., Koshikawa, S., Sagara, H., Miura, T., Yokobori, S. I., Miyagawa, K., Suzuki, Y., Kubo, T., Oyama, M., Kohara, Y., Fujiyama, A., Arakawa, K., Katayama, T., Toyoda, A., & Kunieda, T. (2016). Extremotolerant tardigrade genome and improved radiotolerance of human cultured cells by tardigrade-unique protein. Nature Communications, 7, 12808. https://doi.org/10.1038/ncomms12808

Hashimoto, T. & Kunieda, T. (2017). DNA protection protein, a novel mechanism of radiation tolerance: Lessons from tardigrades. Life (Basel, Switzerland), 7(2), 26. https://doi.org/10.3390/life7020026

Hengherr, S., Brümmer, F., & Schill, R. O. (2008a). Anhydrobiosis in tardigrades and its effects on longevity traits. Journal of Zoology, 275, 216–220. https://doi.org/10.1111/j.1469-7998.2008.00427.x

Hengherr, S., Heyer, A. G., Köhler, H. R., & Schill, R. O. (2008b). Trehalose and anhydrobiosis in tardigrades—evidence for divergence in responses to dehydration. The FEBS Journal, 275(2), 281–288. https://doi.org/10.1111/j.1742-4658.2007.06198.x

Henikoff, S. & Henikoff, J. G. (1992). Amino acid substitution matrices from protein blocks. Proceedings of the National Academy of Sciences, 89, 10915–10919. https://doi.org/10.1073/pnas.89.22.10915

Hesgrove, C., Boothby, T. C. (2020). The biology of tardigrade disordered proteins in extreme stress tolerance. Cell Communication and Signaling, 18(1), 178. https://doi.org/10.1186/s12964-020-00670-2

Hopp, T. P. & Woods, K. R. (1981). Prediction of protein antigenic determinants from amino acid sequences. Proceedings of the National Academy of Sciences, 78(6), 3824–3828. https://doi.org/10.1073/pnas.78.6.3824

Horikawa, D. D., Cumbers, J., Sakakibara, I., Rogoff, D., Leuko, S., Harnoto, R., Arakawa, K., Katayama, T., Kunieda, T., Toyoda, A., Fujiyama, A., & Rothschild, L. J. (2013). Analysis of DNA repair and protection in the tardigrade *Ramazzottius varieornatus* and *Hypsibius dujardini* after exposure to UVC radiation. PLoS One, 8(6), e64793. https://doi.org/10.1371/journal.pone.0064793

Horikawa, D. D., Sakashita, T., Katagiri, C., Watanabe, M., Kikawada, T., Nakahara, Y., Hamada, N., Wada, S., Funayama, T., Higashi, S., Kobayashi, Y., Okuda, T., & Kuwabara, M. (2006). Radiation tolerance in the tardigrade *Milnesium tardigradum*. International Journal of Radiation Biology, 82(12), 843–848. https://doi.org/10.1080/09553000600972956

Hsu, W. L., Oldfield, C., Meng, J., Huang, F., Xue, B., Uversky, V. N., Romero, P., & Dunker, A. K. (2012). Intrinsic protein disorder and protein-protein interactions. Pacific Symposium on Biocomputing, 116–127. https://doi.org/10.1142/9789814366496_0012

Jirgensons, B. (1966). Classification of proteins according to conformation. Die Makromolekulare Chemie, 91, 74–86. https://doi.org/10.1002/macp.1966.020910105

Jones, D. T. & Cozzetto, D. (2015). DISOPRED3: Precise disordered region predictions with annotated protein-binding activity. Bioinformatics, 31(6), 857–863. https://doi.org/10.1093/bioinformatics/btu744

Jönsson, K. I. & Bertolani, R. (2001). Facts and fiction about long-term survival in tardigrades. Journal of Zoology, 255(1), 121–123. https://doi.org/10.1017/S0952836901001169

Jönsson, K. I., Harms-Ringdahl, M., & Torudd, J. (2005). Radiation tolerance in the eutardigrade *Richtersius coronifer*. International Journal of Radiation Biology, 81(9), 649–656. https://doi.org/10.1080/09553000500368453

Jönsson, K. I. & Persson, O. (2010). Trehalose in three species of desiccation tolerant tardigrades. The Open Zoology Journal, 3, 1–5. http://dx.doi.org/10.2174/1874336601003010001

Jönsson, K. I., Rabbow, E., Schill, R. O., Harms-Ringdahl, M., & Rettberg, P. (2008). Tardigrades survive exposure to space in low Earth orbit. Current Biology, 18(17), R729–R731. https://doi.org/10.1016/j.cub.2008.06.048

Jönsson, K. I. & Schill, R. O. (2007). Induction of Hsp70 by desiccation, ionizing radiation and heat-shock in the eutardigrade *Richtersius coronifer*. Comparative Biochemistry and Physiology, 146B, 456–460. https://doi.org/10.1016/j.cbpb.2006.10.111

Jönsson, K. I. (2007). Tardigrades as a potential model organism in space research. Astrobiology, 7(5), 757–66. https://doi.org/10.1089/ast.2006.0088

Jorda, J., Xue, B., Uversky, V. N., & Kajava, A. V. (2010). Protein tandem repeats—the more perfect, the less structured. The FEBS Journal, 277(12), 2673–2682. https://doi.org/10.1111/j.1742-464X.2010.07684.x

Jovine, L., Darie, C. C., Litscher, E. S., & Wassarman, P. M. (2005). Zona pellucida domain proteins. Annual Review of Biochemistry, 74, 83–114. https://doi.org/10.1146/annurev.biochem.74.082803.133039

Kamilari, M., Jørgensen, A., Schiøtt, M, & Møbjerg, N. (2019). Comparative transcriptomics suggest unique molecular adaptations within tardigrade lineages. BMC Genomics, 20, 607. https://doi.org/10.1186/s12864-019-5912-x

Kapp, K., Schrempf, S., Lemberg, M. K., & Dobberstein, B. (2009). Post-targeting functions of signal peptides. In R. Zimmermann (Ed.), Protein Transport into the Endoplasmic Reticulum. Austin (TX): Landes Bioscience. https://www.ncbi.nlm.nih.gov/books/NBK6322/

Kaufmann, J. R., Miller, R., & Cheyne, J. (2011). Vaccine supply chains need to be better funded and strengthened, or lives will be at risk. Health Aff (Millwood), 30(6), 1113–21. https://doi.org/10.1377/hlthaff.2011.0368

Keilin, D. (1959). The Leeuwenhoek lecture—the problem of anabiosis or latent life: History and current concept. Proceedings of the Royal Society B: Biological Sciences, 150, 149–191. https://doi.org/10.1098/rspb.1959.0013

Kinchin, I. M. (2008). Tardigrades and anhydrobiosis water bears and water loss. Biochemistry, 30(4), 18–20. https://doi.org/10.1042/BIO03004018

Kirke, J., Jin, X. L., & Zhang, X. H. (2020). Expression of a tardigrade Dsup gene enhances genome protection in plants. Molecular Biotechnology, 62(11-12), 563–571. https://doi.org/10.1007/s12033-020-00273-9

Kondo, K., Mori, M., Tomita, M., & Arakawa, K. (2020). Pre-treatment with D942, a furancarboxylic acid derivative, increases desiccation tolerance in an anhydrobiotic tardigrade *Hypsibius exemplaris*. FEBS Open Bio, 10(9), 1774–1781. https://doi.org/10.1002/2211-5463.12926

Krendel, M. & Mooseker, M. S. (2005). Myosins: Tails (and heads) of functional diversity. Physiology, 20(4), 239–251. https://doi.org/10.1152/physiol.00014.2005

Kyte, J. & Doolittle, R. F. (1982). A simple method for displaying the hydropathic character of a protein. Journal of Molecular Biology, 157(1), 105–32. https://doi.org/10.1016/0022-2836(82)90515-0

Li, D. & He, X. (2009). Desiccation induced structural alterations in a 66-amino acid fragment of an anhydrobiotic nematode late embryogenesis abundant (LEA) protein. Biomacromolecules, 10, 1469–1477. http://dx.doi.org/10.1021/bm9002688

Liguori, I., Russo, G., Curcio, F., Bulli, G., Aran, L., Della-Morte, D., Gargiulo, G., Testa, G., Cacciatore, F., Bonaduce, D., & Abete, P. (2018). Oxidative stress, aging, and diseases. Clinical Interventions in Aging, 13, 757–772. https://doi.org/10.2147/CIA.S158513

Liu, J., Perumal, N. B., Oldfield, C. J., Su, E. W., Uversky, V. N., & Dunker, A. K. (2006). Intrinsic disorder in transcription factors. Biochemistry, 45, 6873–6888. https://doi.org/10.1021/bi0602718

Luxburg, U. & Schölkopf, B. (2008). Statistical learning theory: Models, concepts, and results. Handbook of the History of Logic, 10, 651–706. https://doi.org/10.1016/B978-0-444-52936-7.50016-1

May, R. M., Maria, M., & Guimard, J. (1964). Actions différentielles des rayons x et ultraviolets sur le tardigrade *Macrobiotus areolatus*, a l’état et desséché. Bulletin biologique de la France et de la Belgique, 98, 349–367.

Michalak, M., Mariani, P., & Opas, M. (1998). Calreticulin, a multifunctional Ca2+ binding chaperone of the endoplasmic reticulum. Biochemistry and Cell Biology, 76(5), 779–785. https://doi.org/10.1139/o98-091

Mínguez-Toral, M., Cuevas-Zuviría, B., Garrido-Arandia, M., & Pacios, L. F. (2020). A computational structural study on the DNA-protecting role of the tardigrade-unique Dsup protein. Scientific Reports, 10. https://doi.org/10.1038/s41598-020-70431-1

Miyazawa, K., Itoh, S. G., Watanabe, H., Uchihashi, T., Yanaka, S., Yagi-Utsumi, M., Kato, K., Arakawa, K., & Okumura, H. (2021). Tardigrade secretory-abundant heat-soluble protein has a flexible β-barrel structure in solution and keeps this structure in dehydration. The Journal of Physical Chemistry B, 125(32), 9145–9154. https://doi.org/10.1021/acs.jpcb.1c04850

Mohan, A., Uversky, V. N., & Radivojac, P. (2009). Influence of sequence changes and environment on intrinsically disordered proteins. PLoS Computational Biology, 5(9), e1000497. https://doi.org/10.1371/journal.pcbi.1000497

Monastyrskyy, B., Kryshtafovych, A., Moult, J., Tramontano, A., & Fidelis, K. (2014). Assessment of protein disorder region predictions in CASP10. Proteins, 2(2), 127–137. https://doi.org/10.1002/prot.24391

NCBI Resource Coordinators. (2016). Database resources of the National Center for Biotechnology Information. Nucleic Acids Research, 44(D1), D7–D19. https://doi.org/10.1093/nar/gkv1290

Necci, M., Piovesan, D., CAID Predictors, DisProt Curators, & Tosatto, S. C. E. (2021). Critical assessment of protein intrinsic disorder prediction. Nature Methods, 18, 472–481. https://doi.org/10.1038/s41592-021-01117-3

Obinata, T., Ono, K., & Ono, S. (2011). Detection of a troponin I-like protein in non-striated muscle of the tardigrades (water bears). Bioarchitecture, 1(2), 96–102. https://doi.org/10.4161/bioa.1.2.16251

Oldfield, C. J. & Dunker, A. K. (2014). Intrinsically disordered proteins and intrinsically disordered protein regions. Annual Review of Biochemistry, 83, 553–584. https://doi.org/10.1146/annurev-biochem-072711-164947

Oldfield, C. J., Meng, J., Yang, J. Y., Yang, M. Q., Uversky, V. N., & Dunker, A. K. (2008). Flexible nets: Disorder and induced fit in the associations of p53 and 14-3-3 with their partners. BMC Genomics, 9 Suppl 1(Suppl 1), S1. https://doi.org/10.1186/1471-2164-9-S1-S1

Parmeggiani, A. (2001). GTPases – molecular switches of cellular signaling pathways. Trends in Biochemical Sciences, 26(1), 76. https://doi.org/10.1016/S0968-0004(00)01713-8

Peng, Z., Mizianty, M. J., Xue, B., Kurgan, L., & Uversky, V. N. (2012). More than just tails: Intrinsic disorder in histone proteins. Molecular BioSystems, 8, 1886–1901. https://doi.org/10.1039/C2MB25102G

Persson, D., Halberg, K. A., Jorgensen, A., Ricci, C., Mobjerg, N., & Kristensen, R. M. (2011). Extreme stress tolerance in tardigrades: Surviving space conditions in low Earth orbit. Journal of Zoological Systematics and Evolutionary Research, 49(Suppl. 1), 90–97. https://doi.org/10.1111/j.1439-0469.2010.00605

Pigoń, A. & Weglarska, B. (1955). Rate of metabolism in tardigrades during active life and anabiosis. Nature, 176, 121–122. https://doi.org/10.1038/176121b0

Piszkiewicz, S., Gunn, K. H., Warmuth, O., Propst, A., Mehta, A., Nguyen, K. H., Kuhlman, E., Guseman, A. J., Stadmiller, S. S., Boothby, T. C., Neher, S. B., & Pielak, G. J. (2019). Protecting activity of desiccated enzymes. Protein Science: A Publication of the Protein Society, 28(5), 941–951. https://doi.org/10.1002/pro.3604

Prasath, T., Greven, H., & D’Haese, J. (2012). EF-hand proteins and the regulation of actin-myosin interaction in the eutardigrade *Hypsibius klebelsbergi* (Tardigrada). Journal of Experimental Zoology Part A: Ecological Genetics and Physiology, 317(5), 311–320. https://doi.org/10.1002/jez.1724

Radivojac, P., Obradovic, Z., Smith, D., Zhu, G., Vucetic, S., Brown, C. J., Lawson, J. D., & Dunker, A. (2004). Protein flexibility and intrinsic disorder. Protein Science, 13, 71–80. https://doi.org/10.1110/ps.03128904

Rahm, P. G. (1921). Biologische und physiologische Beiträge zur Kenntnis der Moosfauna. Zeitschrift für Allgemeine Physiologie, 20, 1–35.

Rebecchi, L., Altiero, T., Guidetti, R., Caselli, V., & Cesari, M. (2011). Resistance of the anhydrobiotic eutardigrade *Paramacrobiotus richtersi* to space flight (LIFE–TARSE mission on FOTON-M3). Journal of Zoological Systematics and Evolutionary Research, 49(Suppl. 1), 1–132. https://doi.org/10.1111/j.1439-0469.2010.00606.x

Rebecchi, L., Altiero, T., Guidetti, R., Cesari, M., Bertolani, R., Negroni, M., & Rizzo, A. M. (2009). Tardigrade resistance to space effects: First results of experiments on the LIFE-TARSE mission on FOTON-M3 (September 2007). Astrobiology, 9(6), 581–591. https://doi.org/10.1089/ast.2008.0305

Rebecchi, L., Altiero, T., Guidetti, R., Cesari, M., Bertolani, R., Rizzo, A. M., & Negroni, M. (2010). Resistance to extreme stresses in the Tardigrada: Experiments on Earth and in space and astrobiological perspectives. Astrobiology Science Conference 2010.

Rebecchi, L. (1997). Ultrastructural study of spermiogenesis and the testicular and spermathecal spermatozoon of the gonochoristic tardigrade *Xerobiotus pseudohufelandi* (Eutardigrada, Macrobiotidae). Journal of Morphology, 234(1), 11–24. https://doi.org/10.1002/(SICI)1097-4687(199710)234:1%3C11::AID-JMOR2%3E3.0.CO;2-Q

Reuner, A., Hengherr, S., Mali, B., Förster, F., Arndt, D., Reinhardt, R., Dandekar, T., Frohme, M., Brümmer, F., & Schill, R. O. (2010). Stress response in tardigrades: Differential gene expression of molecular chaperones. Cell Stress & Chaperones, 15(4), 423–430. https://doi.org/10.1007/s12192-009-0158-1

Ricci, C. & Pagani, M. (1997). Desiccation of *Panagrolaimus rigidus* (Nematoda): Survival, reproduction and the influence on the internal clock. Hydrobiologia, 347, 1–13. https://doi.org/10.1023/A:1002979522816

Richaud, M., Goff, E., Cazevielle, C., Ono, F., Mori, Y., Saini, N., Cuq, P., Baghdiguian, S., Godefroy, N., & Galas, S. (2020). Ultrastructural analysis of the dehydrated tardigrade *Hypsibius exemplaris* unveils an anhydrobiotic-specific architecture. Scientific Reports, 10, 4324. https://doi.org/10.1038/s41598-020-61165-1

Rizzo, A. M., Negroni, M., Altiero, T., Montorfano, G., Corsetto, P., Berselli, P., Berra, B., Guidetti, R., & Rebecchi, L. (2010). Antioxidant defenses in hydrated and desiccated states of the tardigrade *Paramacrobiotus richtersi. Comparative Biochemistry and Physiology*. Part B, Biochemistry & Molecular Biology, 156(2), 115–121. https://doi.org/10.1016/j.cbpb.2010.02.009

Schill, R. O. & Fritz, G. B. (2008). Desiccation tolerance in embryonic stages of the tardigrade *Milnesium tardigradum*. Journal of Zoology (London), 276, 103–107. https://doi.org/10.1111/j.1469-7998.2008.00474.x

Schill, R. O., Steinbrück, G. H. B., & Köhler, H. R. (2004). Stress gene (Hsp70) sequences and quantitative expression in *Milnesium tardigradum* (Tardigrada) during active and cryptobiotic stages. Journal of Experimental Biology, 207, 1607–1613. https://doi.org/10.1242/jeb.0093

Schokraie, E., Hotz-Wagenblatt, A., Warnken, U., Frohme, M., Dandekar, T., Schill, R. O., & Schnölzer, M. (2011). Investigating heat shock proteins of tardigrades in active versus anhydrobiotic state using shotgun proteomics. Journal of Zoological Systematics and Evolutionary Research, 49, 111–119. https://doi.org/10.1111/j.1439-0469.2010.00608.x

Schokraie, E., Hotz-Wagenblatt, A., Warnken, U., Mali, B., Frohme, M., Förster, F., Dandekar, T., Hengherr, S., Schill, R. O., & Schnölzeret, M. (2010). Proteomic analysis of tardigrades: Towards a better understanding of molecular mechanisms by anhydrobiotic organisms. PLoS One, 5(3), e9502. https://doi.org/10.1371/journal.pone.0009502

Schokraie, E., Warnken, U., Hotz-Wagenblatt, A., Grohme, M. A., Hengherr, S., Förster, F., Schill, R. O., Frohme, M., Dandekar, T., & Schnölzer, M. (2012). Comparative proteome analysis of *Milnesium tardigradum* in early embryonic state versus adults in active and anhydrobiotic state. PLoS One, 7(9), e45682. https://doi.org/10.1371/journal.pone.0045682

Seki, K. & Toyoshima, M. (1998). Preserving tardigrades under pressure. Nature, 395, 853–854. https://doi.org/10.1038/27576

Simon, M. & Hancock, J. M. (2009). Tandem and cryptic amino acid repeats accumulate in disordered regions of proteins. Genome Biology, 10(6), R59. https://doi.org/10.1186/gb-2009-10-6-r59

Sloan, D., Alves Batista, R., & Loeb, A. (2017). The resilience of life to astrophysical events. Scientific Reports, 7, 5419, https://doi.org/10.1038/s41598-017-05796-x

Smathers, R. L. & Petersen, D. R. (2011). The human fatty acid-binding protein family: Evolutionary divergences and functions. Human Genomics, 5(3), 170–191. https://doi.org/10.1186/1479-7364-5-3-170

Sun, W. & Leopold, A. (1997). Cytoplasmic vitrification and survival of anhydrobiotic organisms. Comparative Biochemistry and Physiology Part A: Physiology, 117(3), 327–333. https://doi.org/10.1016/S0300-9629(96)00271-X

Tanaka, S., Tanaka, J., Miwa, Y., Horikawa, D. D., Katayama, T., Arakawa, K., Toyoda, A., Kubo, T., & Kunieda, T. (2015). Novel mitochondria-targeted heat-soluble proteins identified in the anhydrobiotic tardigrade improve osmotic tolerance of human cells. PLoS One, 10(2), e0118272. https://doi.org/10.1371/journal.pone.0118272

Tardigrade [Fact sheet]. (n.d.). National Geographic. Retrieved September 5, 2021, from https://www.nationalgeographic.com/animals/invertebrates/facts/tardigrades-water-bears

Tsujimoto, M., Imura, S., & Kanda, H. (2016). Recovery and reproduction of an Antarctic tardigrade retrieved from a moss sample frozen for over 30 years. Cryobiology, 72(1), 78–81. https://doi.org/10.1016/j.cryobiol.2015.12.003

UniProt. (2021). The UniProt Consortium, UniProt: The universal protein knowledgebase in 2021. Nucleic Acids Research, 49(D1), D480–D489, https://doi.org/10.1093/nar/gkaa1100

Uversky, V. N. & Dunker, A. K. (2010). Understanding protein non-folding. Biochimica et Biophysica Acta, 1804, 1231–1264. https://doi.org/10.1016/j.bbapap.2010.01.017

Vaishnavi, R. & Lorsch, J. R. (2012). Chapter twenty-five—ATP and GTP hydrolysis assays (TLC). Methods in Enzymology, 533, 325–334. https://doi.org/10.1016/B978-0-12-420067-8.00025-8

Van der Lee, R., Buljan, M., Lang, B., Weatheritt, R. J., Daughdrill, G. W., Dunker, A. K., Fuxreiter, M., Gough, J., Gsponer, J., Jones, D. T., Kim, P. M., Kriwacki, R. W., Oldfield, C. J., Pappu, R. V., Tompa, P., Uversky, V. N., Wright, P. E., & Babu, M. M. (2014). Classification of intrinsically disordered regions and proteins. Chemical Reviews, 114(13), 6589–6631. https://doi.org/10.1021/cr400525m

Vukich, M. Ganga, P. L., Cavalieri, D., Rizzetto, L., Rivero, D., Pollastri, S., Mugnai, S., Mancuso, S., Pastorelli, S., Lambreva, M., Antonacci, A., Margonelli, A., Bertalan, I., Johanningmeier, U., Giardi, M. T., Rea, G., Pugliese, M., Quarto, M., Roca, V., Zanini, A., Borla, O., Rebecchi, L., Altiero, T., Guidetti, R., Cesari, M., Marchioro, T., Bertolani, R., Pace, E,., De Sio, A., Casarosa, M., Tozzetti, L., Branciamore, S., Gallori, E., Scarigella, M., Bruzzi, M., Bucciolini, M., Talamonti, C., Donati, A., & Zolesi, V. (2012). BIOKIS: A model payload for multidisciplinary experiments in microgravity. Microgravity Science and Technology, 24, 397–409. https://doi.org/10.1007/s12217-012-9309-6

Wang, C., Grohme, M. A., Mali, B., Schill, R. O., & Frohme, M. (2014). Towards decrypting cryptobiosis—analyzing anhydrobiosis in the tardigrade *Milnesium tardigradum* using transcriptome sequencing. PloS One, 9(3), e92663. https://doi.org/10.1371/journal.pone.0092663

Webb, S. J. (1964). Bound water, metabolites and genetic continuity. Nature, 203, 374–377. https://doi.org/10.1038/203374a0

Westover, C., Najjar, D., Meydan, C., Grigorev, K., Veling, M. T., Iosim, S., Colon, R., Yang, S., Restrepo, U., Chin, C., Butler, D., Moszary, C., Rahmatulloev, S., Afshinnekoo, E., Silver, P. A., Chang, R. L., & Mason, C. E. (2020). Engineering radioprotective human cells using the tardigrade damage suppressor protein, DSUP. BioRxiv. https://doi.org/10.1101/2020.11.10.373571

Wright, P. E. & Dyson, H. J. (2015). Intrinsically disordered proteins in cellular signaling and regulation. Nature Reviews, Molecular Cell Biology, 16(1), 18–29. https://doi.org/10.1038/nrm3920

Yamaguchi, A., Tanaka, S., Yamaguchi, S., Kuwahara, H., Takamura, C., Imajoh-Ohmi, S., Horikawa, D. D., Toyoda, A., Katayama, T., Arakawa, K., Fujiyama, A., Kubo, T., & Kunieda, T. (2012). Two novel heat-soluble protein families abundantly expressed in an anhydrobiotic tardigrade. PLoS One, 7(8), e44209. https://doi.org/10.1371/journal.pone.0044209

Yin, L. M., Schnoor, M., & Jun, C. D. (2020). Structural characteristics, binding partners and related diseases of the calponin homology (CH) domain. Frontiers in Cell and Developmental Biology, 8, 342. https://doi.org/10.3389/fcell.2020.00342

Yoshida, Y., Koutsovoulos, G., Laetsch, D. R., Stevens, L., Kumar, S., Horikawa, D. D., Ishino, K., Komine, S., Kunieda, T., Tomita, M., Blaxter, M., & Arakawa, K. (2017). Comparative genomics of the tardigrades *Hypsibius dujardini* and *Ramazzottius varieornatus*. PLoS Biology, 15(7), e2002266. https://doi.org/10.1371/journal.pbio.2002266

Zaffran, M., Vandelaer, J., Kristensen, D., Melgaard, B., Yadav, P., Antwi-Agyei, K. O., & Lasher, H. (2013). The imperative for stronger vaccine supply and logistics systems. Vaccine, 31(2), B73–B80. https://doi.org/10.1016/j.vaccine.2012.11.03

